# Listening to tropical forest soils

**DOI:** 10.1101/2023.05.19.541323

**Authors:** Oliver Metcalf, Fabricio Baccaro, Jos Barlow, Erika Berenguer, Tom Bradfer-Lawrence, Liana Chesini Rossi, Érica Marinho do Vale, Alexander Lees

**Affiliations:** Division of Biology and Conservation Ecology, Department of Natural Sciences, Manchester Metropolitan University, Manchester, UK; Departamento de Biologia, Universidade Federal do Amazonas, Manaus, Brazil; Lancaster Environment Centre, Lancaster University, Lancaster, Lancashire, UK; Environmental Change Institute, University of Oxford, Oxford OX1 3QY, UK; Centre for Conservation Science, RSPB, 2 Lochside Drive, Edinburgh, UK; Biological and Environmental Sciences, University of Stirling, Stirling, UK; Rio de Janeiro Botanical Garden, Rio de Janeiro, RJ, Brazil; The National Institute of Amazonian Research (INPA), Manaus, (Amazonas), Brazil

## Abstract

PAM has proven to be an effective tool for monitoring biotic soundscapes in the marine, terrestrial, and aquatic realms. Recently it has been suggested that it could also be an effective method for monitoring soil fauna, but has so far been used in only four studies in temperate and polar regions. We present the first study of soil soundscapes in tropical forests, using a novel analytical pipeline allowing for the use of in-situ recording of soundscapes with minimal soil disturbance. We found significant differences in soil soundscapes between burnt and unburnt forests and the first indications of a diel cycle in soil soundscapes. These promising results and methodological advances highlight the potential of PAM for large-scale and long-term monitoring of soil biodiversity. We use the results to discuss research priorities, including relating soil biophony to biodiversity, community structure and ecosystem functioning, and the use of appropriate hardware and analytical techniques.

## Introduction

Passive acoustic monitoring (PAM) has allowed the field of ecoacoustics to reveal new information on a range of soniferous communities in terrestrial above-ground and aquatic habitats (Desjonquères et al., 2020, Sugai et al., 2019). In particular, it has proven effective at showing that soundscapes vary significantly across landscapes and habitat types at large spatial scales (Mitchell et al., 2021, Metcalf et al., 2021, Bradfer-Lawrence et al., Eldridge et al., Do Nascimento et al., 2020), and in revealing temporal dynamics such as diel and seasonal variation in a range of habitats from tropical forests to coral reefs (Garcia Oliveira et al., 2021, Bertucci et al., 2016).

PAM has recently been suggested as a tool for studying soil soundscapes (Maeder et al., 2019). Soil faunas are highly diverse and functionally important (Bardgett et al., 2014), playing important roles in nutrient cycling, carbon sequestration and soil structure (Briones, 2014; Lavelle et al., 2006). However, studying soil fauna is challenging; existing methods are labour intensive, time consuming, often invasive, and require specialist taxonomic knowledge (Geisen et al., 2019). Consequently there are some significant gaps in the soil ecology literature, with knowledge particularly lacking for studies covering large spatial extents and temporal dynamics (Guerra et al., 2020) - areas PAM may be particularly well-suited to resolving.

Despite being apparently well-suited to the task, ecoacoustic analytical techniques have rarely been used to monitor soundscape dynamics below the ground. Only four studies have used soundscapes to compare soil biodiversity in different locations (Keen et al.; 2022, Maeder et al.; 2019, Maeder et al., 2022; Robinson et al., 2023), all of which used different methodologies and equipment, and have all been conducted in temperate and polar regions. We are unaware of any studies from the tropics. Monitoring changes in soil biodiversity is of particular importance in tropical forests, as these are amongst the most diverse habitats in the world, and amongst the most threatened by human impact (Barlow et al., 2018). Anthropological impacts such as forest disturbance have already been shown to negatively impact soil fauna (Franco et al., 2019), although the temporal and spatial dimensions of such impacts are poorly understood.

As soil soundscape assessment is a novel application of ecoacoustics in the tropics, it is first necessary to make basic assessments of the capacity to differentiate soundscapes from different habitats and time of day, prior to application at large spatial and temporal scales. Here we develop a novel pipeline for processing acoustic data to assess soil soundscapes in tropical rainforests. This application is particularly pertinent as forest disturbance, such as edge effects, selective logging, wildfire, and increasing drought frequency (Lapola et al., 2023) is pervasive across Amazonia, and understanding its impact is considered a conservation priority. We test the method’s suitability by comparing soundscapes from burnt and unburnt forests, and by assessing biophonic activity patterns across the diel cycle. To do so, we test three hypotheses: Hypothesis 1 “detecting biophony”: soil recordings have non-random structure indicative of biophony - sounds generated from biological sources; Hypothesis 2 “sensitivity to disturbance”: Soil soundscapes are sensitive to forest disturbance with unburnt forest soundscapes statistically differentiated from burnt forest soundscapes; and Hypothesis 3 “temporal periodicity”: Soil soundscapes exhibit temporal periodicity.

## Methods

### Data Collection

We collected two soil acoustic datasets in three municipalities: Santarém, Belterra and Mojuí dos Campos, in the state of Pará, (eastern Brazilian Amazon latitude ∼ −3.046, longitude −54.947 WGS 84). The region has a hot and humid climate with a marked dry spell between August and November, and annual average temperatures of 25°C, 86% mean relative humidity and a mean 1920 mm of rain (Berenguer et al., 2018). In general, soils are rich in clay and nutrient poor (Silver et al., 2000). All recordings were made in *terra firme* forests - i.e. those not seasonally flooded. For full details on the study region, see Gardner et al. (2013).

To test whether soil recordings include biophony and are sensitive to forest disturbance, we sampled seven sites (hereafter ‘spatial dataset’) unaffected by fire and three sites in which the forest has been burnt during the prolonged drought associated with the El Niño events of 2015/2016 (Withey et al., 2018). All sites were separated by a minimum distance of 2⍰km. Data were collected between 21 November and 06 December 2023. This period is the onset of the rainy season when soil biotic activity is likely at its highest (Levings and Windsor, 1985) - although we avoided recording during periods of rain.

Recordings were made at each point for 30 minutes between 09:20 and 14:00, with a minimum buffer of 3 minutes at the start and end of each recording to avoid including footsteps or other anthropogenic sounds associated with researcher presence. Recordings were made using a Zoom H5n recorder with JrF C-series Pro contact microphones and XLR impedance adapters in both the left and right channels. The input levels were set to the maximum (10). We chose to use contact microphones, as they are much less sensitive to above-ground sounds, and therefore reducing the likelihood of recording above ground sounds that may obscure the soil soundscape pattern.

Microphones were placed at the furthest distance apart the cables would allow, approximately 5 m, and small holes were dug so that the microphones would sit in the soil/clay layer just below the hummus layer (see SOM Appendix 1 for video of standard deployment and SOM Appendix 2 for an analysis of the independence of each channel). This method of microphone deployment thus only caused a minimal amount of soil disturbance, ensuring that recordings were likely to be as unimpacted by the experimental setup as possible.

To test whether soundscapes exhibit temporal periodicity (hereafter “temporal” dataset), we sampled three new sites located a minimum of 100 m apart. Recording was conducted using the equipment set-up described above for 24 hours on 7^th^ December 2022 at Site 1, for 48 hours starting on 8^th^ December 2022 at Site 2 and 24 hours on 12^th^ December 2022 at Site 3.

### Analysis

We split all recordings from both datasets into one minute files. We tested whether the recordings contained biophony with a qualitative analysis conducted by viewing a proportion of the spatial dataset in Raven Pro (ver 1.6; Center for Conservation Bioacoustics, 2019). We compared the recordings to data from the Sounding Soil repository (https://www.soundingsoil.ch/en/listen/) until the reviewers (OCM, ACL) were happy that biophony was evident. It was clear that the vast majority of sound was at frequencies <1 kHz, and often much lower. Consequently, we used a macro in Audacity (ver 2.3.3; Audacity team, 2021) to shift the pitch upwards by 900%, such that sounds in the original data at 1 kHz were now at 10 kHz. In the text hereafter, we refer to the frequencies at their original values. To remove prominent non-biophonic noise from the recordings (most likely microphone self-noise), we applied an adaptive level equalisation algorithm (Towsey, 2013) using the remove_background_along_axis function in the scikit-maad package (ver 1.3.12.; Ulloa et al., 2021) in Python (Van Rossum & Drake, 2009).

To test the sensitivity of soil soundscapes to disturbance, we calculated a suite of acoustic indices to give statistical summaries of the soundscape. Given the general paucity of information on the dynamics of soil biophony, we selected six acoustic indices that reflect a range of soundscape patterns; the Acoustic Complexity Index (ACI, Pieretti et al., 2011), the Bioacoustic Index (BI, Boelman et al., 2008), the number of spectral events per second (EVNspCount, Towsey et al., 2013, QUT, 2023), an adjusted version of the Normalized-Difference Sound Index (NDSI, Kasten et al., 2012), the proportion of the spectrogram covered by regions of interest (ROIcover, Ulloa et al., 2021) and Frequency Entropy (Hf, Sueur et al., 2008) - see SOM Appendix 3 for full details of the index parameters and our expectation of their responses to increased biotic soil activity. We calculated each of the acoustic indices at 0-500 Hz. To assess the sensitivity of acoustic index values to the frequency parameters and denoising techniques selected, we tested the impact of two different noise removal techniques and three different frequency bands to calculate the indices, the results of which are available in SOM Appendix 4. All acoustic index values were scaled between zero and one prior to analysis.

We compared the scores from unburnt and burnt sites using a generalised linear mixed model for each index. We hypothesised that soil biotic activity would decline with disturbance, represented by lower ACI, BI, EVNspCount, Hf and ROI cover and higher NDSI values in burnt forest. We used the glmmTMB R package (ver 1.1.5, Brooks et al., 2017, R Core Team, 2022) using the same structure for each model, with the acoustic index as the dependent variable, the burnt/unburnt forest class as a two-level independent variable, and recording location and recording channel as a nested random factor with a beta family distribution using a logit link. The beta distribution has been used for analysing acoustic indices previously (Bradfer-Lawrence et al., 2020). As values range from 0 to 1 in the beta distribution it is well suited to normalised data and has no prior expectations related to the distribution within that range, so can handle heteroskedastic and asymmetrical data (Ferrari and Cribari-Neto, 2004).

Finally, to assess temporal periodicity in soil soundscapes, we looked for the presence of a diel cycle in soil acoustic activity. Using the temporal dataset, we computed indices in the same manner as above. We used hierarchical general additive models following Pedersen et al., (2019) to assess temporal patterns in the index values. Again, we retained the same model structure for each model, with index value as the dependent variable, a smooth term for time as an independent variable with a cyclic cubic regression spline and six basis dimensions, and a factorial smooth of recording channel by recording site with a random effect spline. We used a beta family distribution with logit link and restricted maximum likelihood estimation for the smoothing parameters.

## Results

Qualitative assessment of the sound recordings suggested clear patterns that appeared consistent with biophony in soil soundscapes from other regions, providing strong support for detection of biophony in our recordings. Biophony appeared to be present and often dominant in many of the recordings. Biophonic sounds generally consisted of ‘knocking’ and ‘scratching’ noises (Figure 1), corresponding well to the recordings of soil biophony in the Sounding Soil repository. However, inspection of spectrograms across the human audible frequency ranges (e.g. 0-20 kHz, Purves et al., 2001) often used to assess above-ground soundscape recordings were not useful. When listening to the recordings at their original frequencies it was difficult to distinguish biophony from microphone self-noise and environmental sound, suggesting that a degree of sound processing was required. The pitch-shifting process in Audacity enhanced the clarity of these sounds, revealing possible evidence of alternating signalling (Figure 1). An example recording with this alternating signalling is available on Dryad, as is the full spatial dataset (see the Data Availability statement)

**Figure 1.**
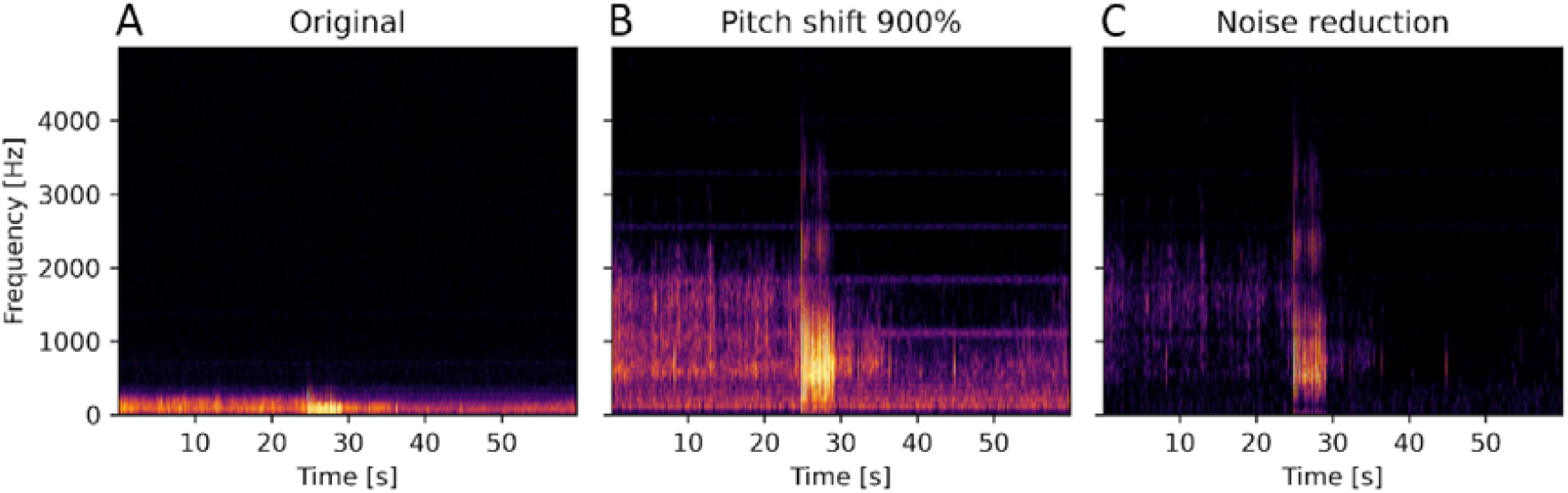
Spectrograms of one minute of audio recording from a burnt forest site on 21^st^ November 2022 at 09:47 in its untransformed state (A), after applying a 900% pitch-shift transformation in Audacity (B) and following denoising (C). The sound event at approximately 25 s appears to be biophonic in origin.

We also found support for our hypothesis that soil biophony changes following forest disturbance with significant differences (p =<0.05) in the values of BI and EVNspCount in burnt and unburnt forest (Figure 2). However, these indices responded differently to disturbance than anticipated, with increased values in burnt forest (mean BI in burnt forest 0.31±0.12 (SD), unburnt forest 0.21±0.10 (SD); mean EVNspCount in burnt forest 0.12±0.13 (SD); unburnt forest 0.07±0.10(SD)). ACI, Hf, NDSI, and ROIcover were not significant.

**Figure 2.**
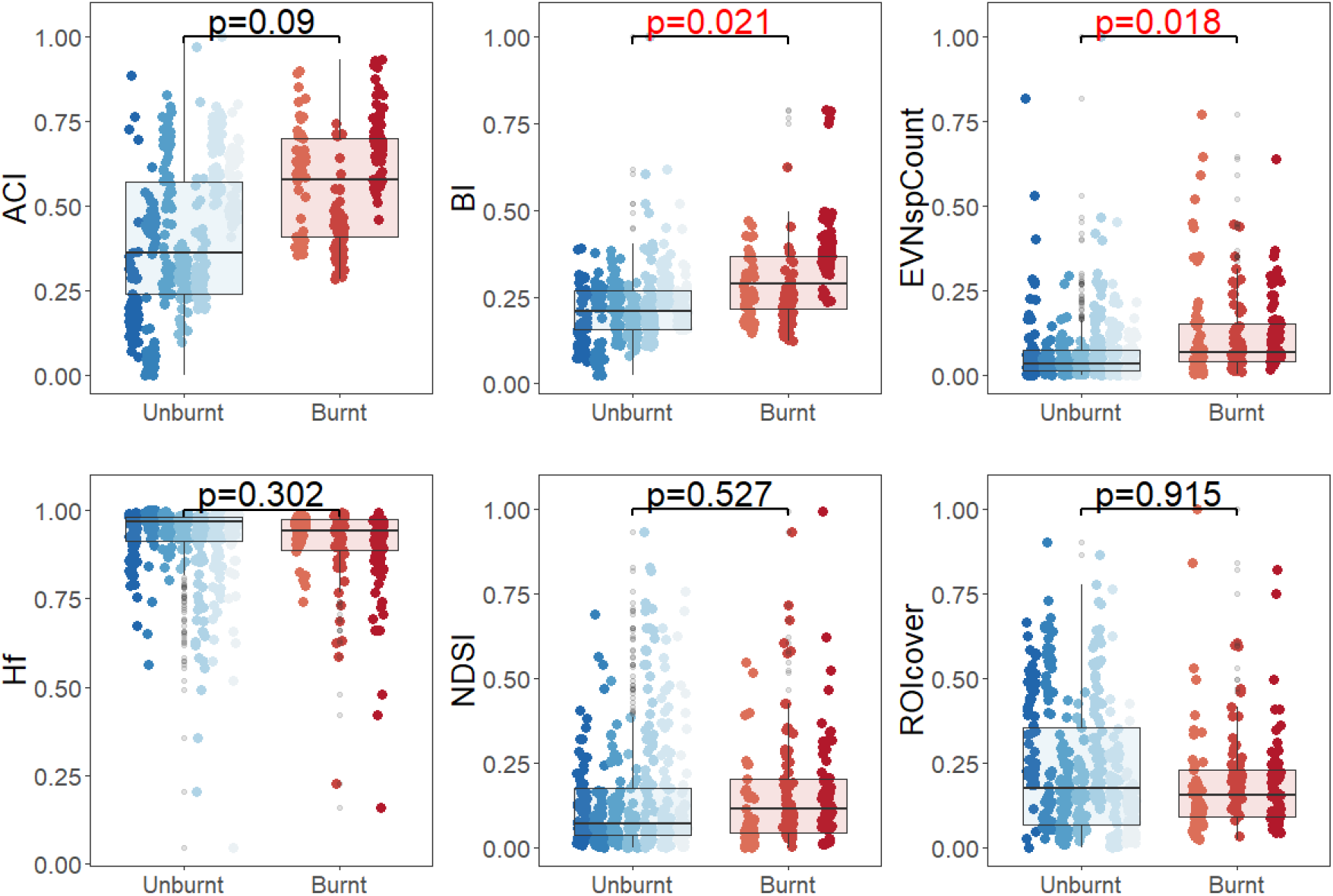
Boxplots of the acoustic index values for six acoustic indices. Points are jittered by recording site. Significance values are derived from generalised linear mixed models with red text indicating p < 0.05.

Analysis of temporal periodicity demonstrated variation in the values of ACI, BI, and EVNspCount and ROIcover (Figure 3), with peaks between 08:00 and 12:00, and lows between 23:00 and 02:00, providing strong evidence for diel cycles in soil acoustic activity. Hf showed an inverse trend, with low values just before 08:00 and high values around midnight, whilst NDSI showed two peaks, one at around 04:00 and the second at 16:00, with lows at around 10:00 and 20:00.

**Figure 3.**
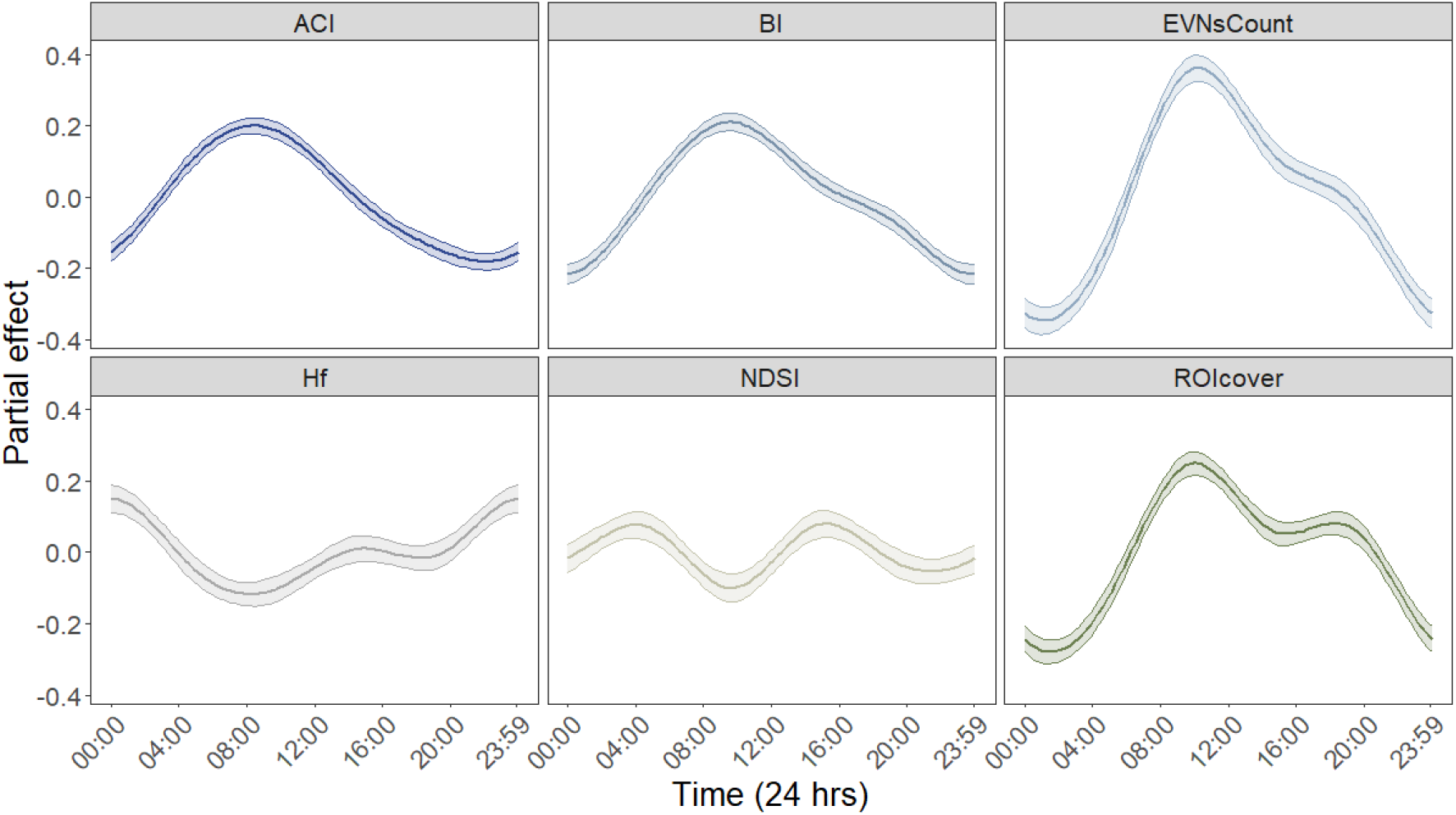
The effect of time of day on six acoustic indices.

## Discussion

### Methodological considerations

We present strong evidence that PAM can be a valuable tool for monitoring tropical forest soil soundscapes. We found that the soil soundscape is dominated by much lower frequencies than those above ground, meaning that some form of pre-processing of the sound is beneficial in order to use standard ecoacoustic analysis techniques such as indices. In this case, we chose to use pitch-shifting and denoising techniques. Still, there are a range of alternatives that may be as effective, such as customising the acoustic index calculations for lower frequencies and using more expensive equipment that can record with less recorder self-noise.

We identified a range of biotic sounds, but assigning them to any taxonomic group is currently not possible as there are very few, if any, reference sounds available for tropical soil fauna. Creating a reference library of soil biotic sound should be a research priority for this ecoacoustic field and would greatly facilitate understanding the composition of the soil soundscape. Although it may be quite some time, if ever, before soil taxa can be inventoried as effectively as birds with PAM recordings. Rather, soil biophony may be best utilised to generate community level metrics, perhaps indicative of macrofaunal biomass or representative of ecosystem functions, such as bioturbation rates.

### Soundscape patterns

We were also able to distinguish between soil soundscapes from burnt and unburnt forests using acoustic indices. This capacity to discriminate between habitats by soundscape is in keeping with above ground ecoacoustic analysis (Bradfer-Lawrence et al., 2019, Do Nascimento et al., 2020, Metcalf et al., 2021). Although the trend in index values was the inverse to that hypothesised, we should be cautious in ascribing this to an increase in biodiversity in burnt forest. A more likely scenario may be the competitive release of one or two generalist or fire-resilient species and subsequent increase in abundance causing the increase in BI and EVNspCount values we found. However, it is very difficult to infer direct biodiversity change from acoustic index values, and it is not clear how effective acoustic indices are as proxies for traditional biodiversity metrics. There are a large number of studies linking above-ground acoustic indices to biodiversity metrics such as species richness, diversity, or abundance, but none of the acoustic indices have a clear and consistent relationship with these metrics (Alcocer et al., 2022), even for well-studied taxa like birds. Whilst attempting to link soil soundscapes directly to diversity metrics is obviously desirable, and linking acoustic indices from soil soundscapes to more traditional measures of soil diversity such as those derived from pit-traps should still be considered a research priority - it may be better to link soundscapes to acoustic morphotypes or ‘operational sound units’ (Luypaert et al., 2022), or to soil functioning such as bioturbation. This may be especially true for soil functions associated with a few abundant species, especially if they produce sound or the functional processes themselves produce sound.

We believe our study is also the first acoustic evidence for diel cycles in soil fauna in the tropics, and there is very limited research on the topic using any method. Our findings support similar results from Switzerland (Maeder et al., 2022), showing strong daytime peaks in acoustic activity. Interestingly the Swiss study finds peaks in ACI values after midday, in line with maximum surface temperature. In contrast, our results show ACI peaking just before 09:00, well before the average surface temperature peaks at our study site (13:00 during the survey period, calculated using the weathermetrics package in R (Anderson et al, 2013). Previous studies have shown roughly similar peaks in soil insect activity using direct measurements such as trapping (Williams 1959), but studies into the diel cycle or circadian rhythms of soil fauna are rare. Further research is required to explore whether this diel cycle is universal amongst soil fauna or, as seems more likely, there are multiple trends, as in birds where some species avoid vocalising at the peak of the dawn chorus (Metcalf et al., 2021), and our method was only sensitive enough to detect the dominant pattern.

### Summary

We show that PAM can facilitate research into important and understudied aspects of soil biodiversity (Guerra et al., 2020) and in regions of high conservation value. Substantially more research is required to refine ecoacoustics as a tool for monitoring soil fauna. Still, given the early stage of development of the method, our results demonstrate the potential value of PAM for soil research in the tropics, producing novel insights into soil biotic activity within just a few months of data collection.

## Acknowledgements

We would like to thank the RAS field and laboratory assistants: Marcos Oliveira, Gilson Oliveira, Renílson Freitas, and Josué Jesus de Oliveira for their hard work and assistance, without whom this would not be possible. We are also grateful to Joice Ferreira for logistical field support in Brazil.

Fieldwork in Brazil and later analysis was supported by research grants PELD-RAS (CNPq/CAPES/PELD 441659/2016-0) and the BNP Paribas Foundation’s Climate and Biodiversity Initiative (Project Bioclimate).

## Authorship statement

All authors made substantial contributions. Oliver Metcalf, Jos Barlow, Alexander Lees, Fabricio Baccaro and Tom Bradfer-Lawrence made substantial contributions to conception and design. Oliver Metcalf, Erika Berenguer, Liana Chessini Rossi and, Érica Marinho do Vale contributed to the acquisition of data. All authors contributed to analysis and interpretation of data, and drafting the article. All authors had final approval of the version to be published, agree to be accountable for the aspects of the work that they conducted and ensuring that questions related to the accuracy or integrity of any part of their work are appropriately investigated and resolved.

## Data Archiving statement

The spatial dataset is available on Dryad.

## SOM

## Appendix 1: Video of recording unit deployment at one of the spatial survey locations

## Appendix 2: A comparison of independence between recording channels.

As soil ecoacoustics is a new field, there was little information available to us on the likely spatial scale of soil communities, or detection distances for the contact microphones under the soil. Although we believe we have largely addressed spatial autocorrelation in our models by including the recording channel as a random effect in both analyses, we felt it would be beneficial to include some supplementary analysis of the overlap in sound detection between the microphones.

Firstly, we conducted qualitative analysis by creating long-duration spectrograms of the 24 hr period from Site 1 of the temporal dataset (Fig S2.1), showing the first 10 s of each minute over the 24 hr period. It is also worth noting that the footsteps of departing and returning researchers were audible on both channels in the discarded portions at the start and end of all recordings.

**Figure S2.1.**
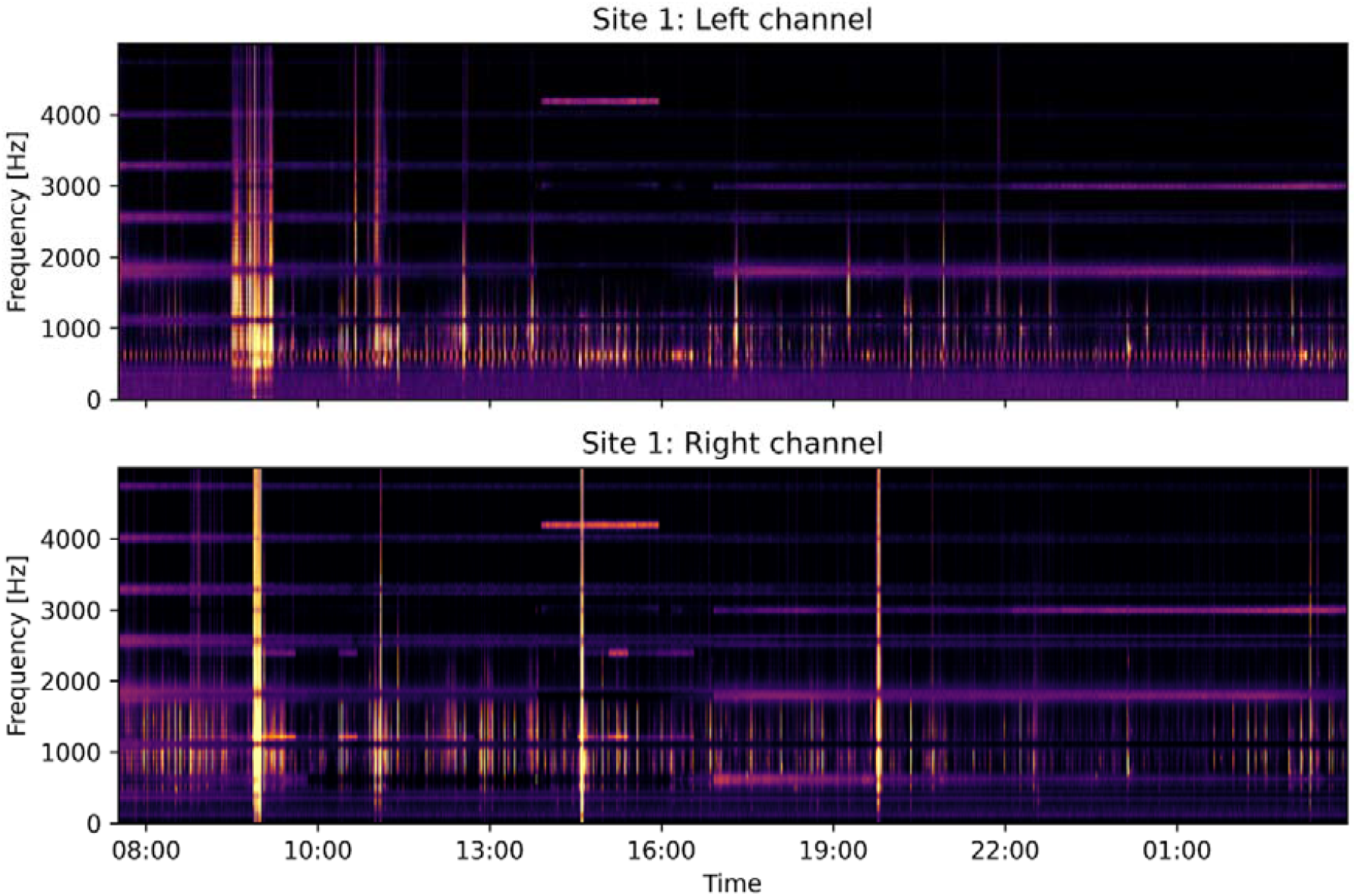
Long duration spectrograms of the 24 hr period from Site 1 of the spatial dataset, comparing the left channel (top) with the right channel (bottom). Whilst there is some similarity in trends between channels, in general there is limited direct correspondence between individual sound events, suggesting a high degree of independence between the channels.

Secondly, we undertook a quantitative analysis using the spatial dataset. We calculated the Spearman’s Rank correlation between the index values from the left and right channels of each site (same-site correlations) using the cor package in R. Next we calculated the pairwise correlation of the left channel of a site with the left channel from each remaining site, and did the same for the right channels (cross-site correlations). Finally we used a generalised linear mixed effect model to compare the same-site correlations with the cross-site correlations to see if index values were more strongly correlated if they came from the left and right channels of the same site. We used the glmmTMB R package with the correlation coefficient as the dependent variable and a two-level independent variable of same-site and cross-site correlation, with acoustic index as a random factor and a beta family distribution using a logit link. Due to the beta family distribution, to make the correlation coefficients fit between the 0-1 scale, we added 0.00001 to the two values that had a correlation coefficient of 0.

We found that there appeared to be limited correspondence between sounds in the long-duration spectrogram, implying a high degree of independence between the channels. In the second analysis, correlation between the index values from the left and right channel recordings at the same site was low - ACI had the highest mean correlation of 0.26±0.24 (SD) and NDSI had the lowest 0.11±0.04. Overall, there was no significant difference (p=0.903) in the correlation coefficients for same-site correlation as there was for crossed-site correlation (Fig SOM2.2). Such low correlations imply that there is minimal overlap in sound events in soil soundscapes over even very short distances (∼ 5 m), implying a high level of local spatial variation.

**Figure S2.2.**
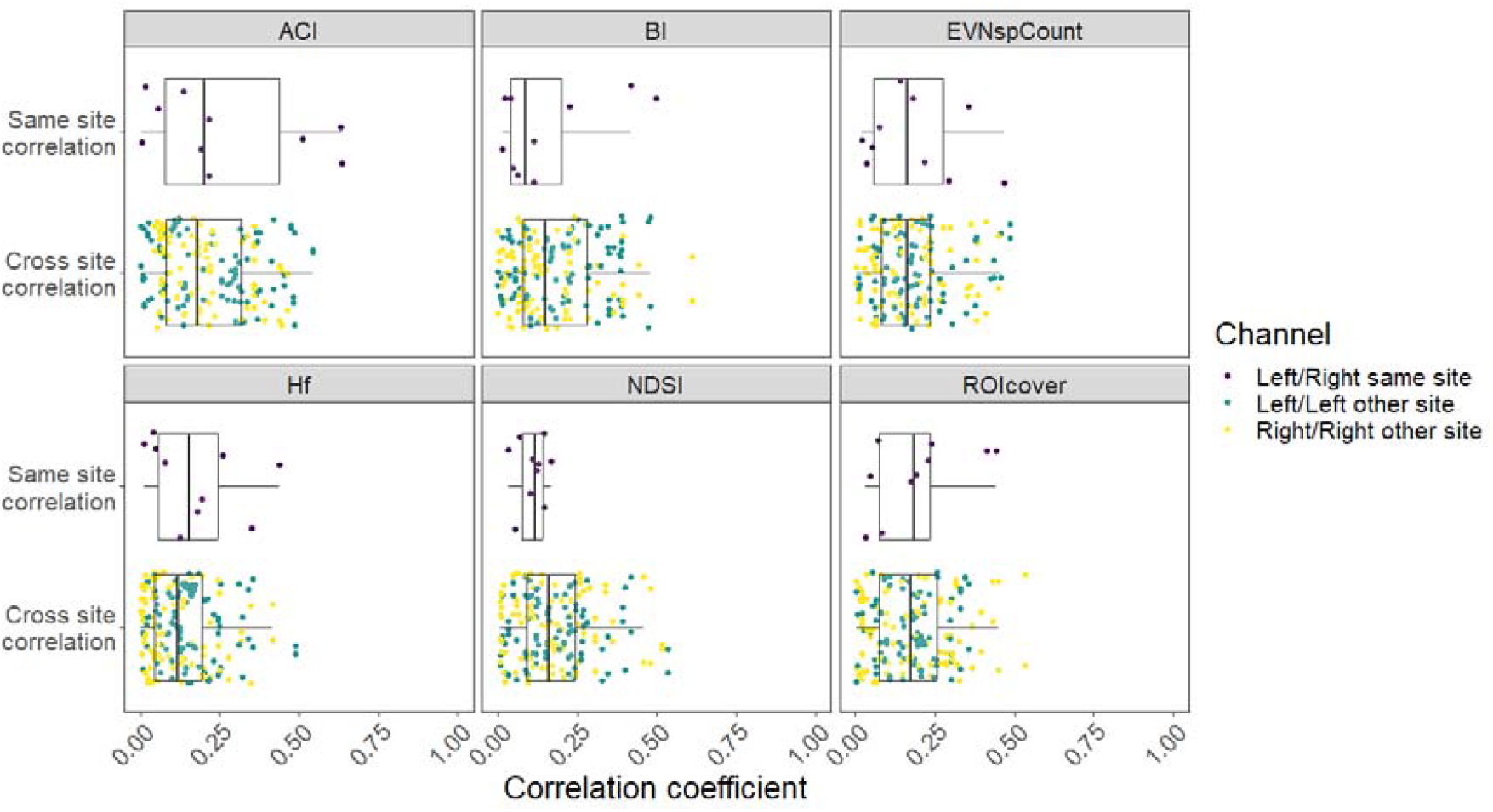
Spearman’s rank correlation coefficient values comparing correlation of acoustic index values derived from both channels at the same site (Same site correlation), with correlation values from the corresponding channels at other sites (Cross site correlation). Overall, correlation was low, and same site correlations were not significantly larger than cross site correlations, indicating a high degree of independence between channels from the same site and potentially a high level of local soundscape variation between sites.

## SOM Appendix 3: Acoustic Index calculation parameters, descriptions, and hypotheses.

We computed six indices (Table S1): the Acoustic Complexity Index (ACI) at default settings; the Bioacoustic Index (BI), starting at 10 Hz and up to the maximum of each bandwidth using the ‘soundecology’ method; the number of spectral events per second (EVNspCount), with a 6-db threshold and a minimum duration of 0.2 s; Normalized-Difference Soundscape Index (NDSI), with the lower frequency band set at 0 to 150 and Hz and upper frequency band set at 150 Hz to 500 Hz; and the proportion of the spectrogram covered by regions of interest (ROIcover), with the default amplitude settings, a max xy ratio of 10, a mask 1 parameter of 6 and a mask 2 parameter of 0.2. Finally we calculated Temporal Entropy (Hf) using the default settings and with compatibility set to the seewave package. Full details of these settings can be found in Ulloa et al (2021).

**Table S3.1.**
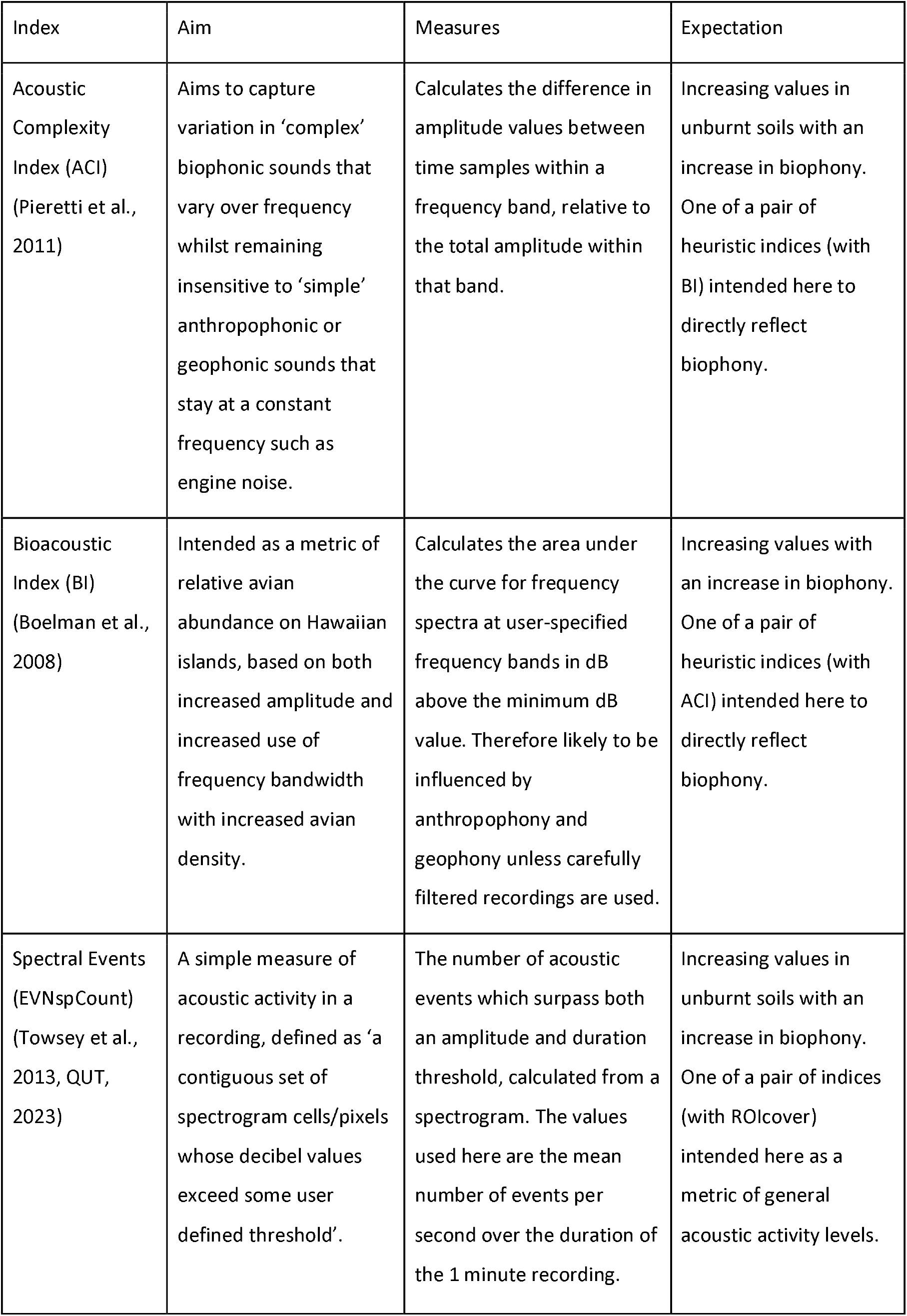
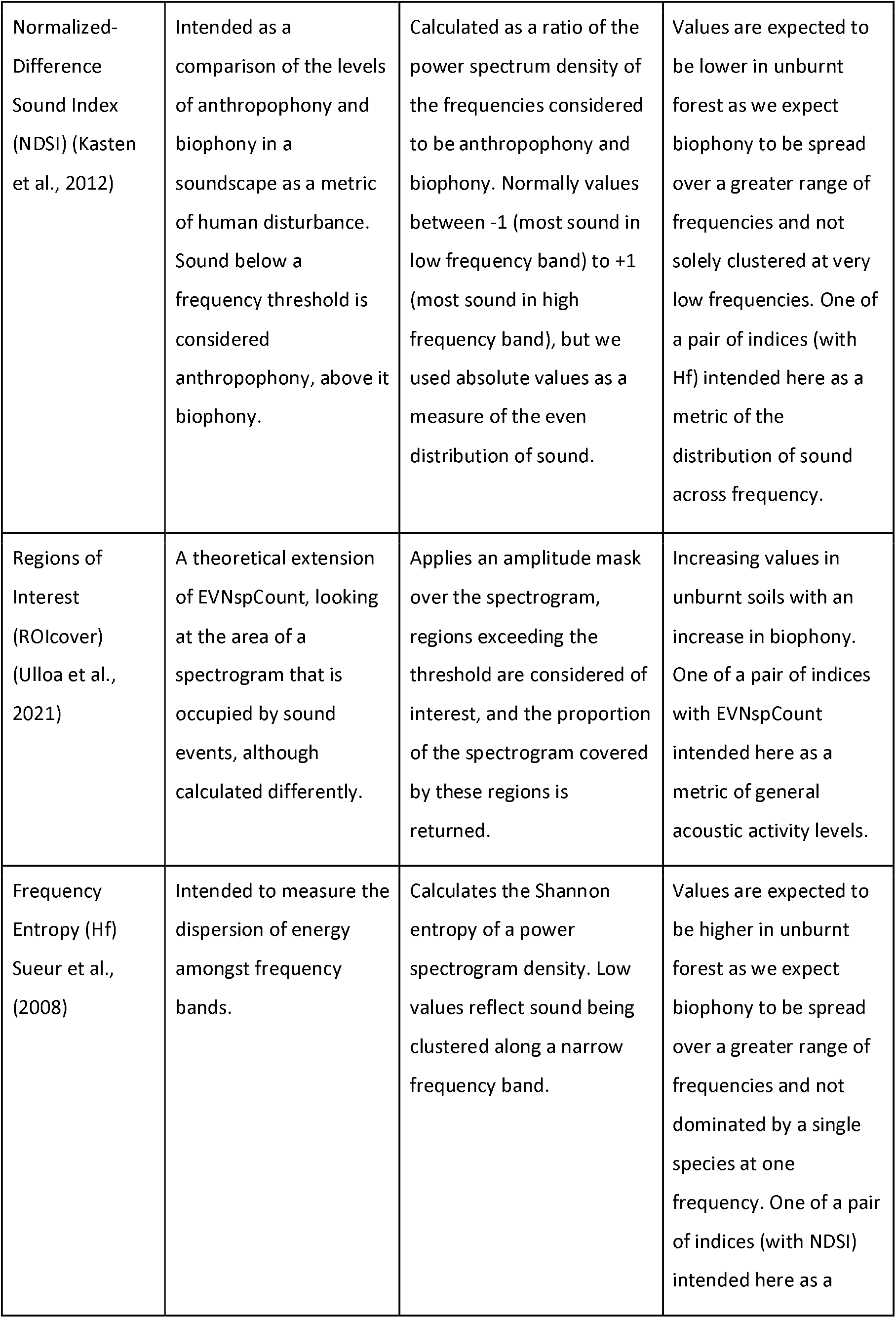
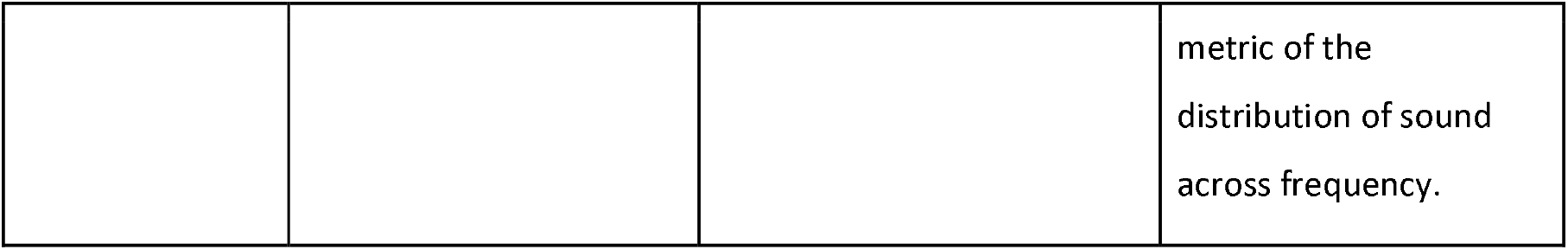
Acoustic indices used in this study, showing the patterns the index is designed to capture, the way it measures this, and our expectations of the way these would be reflected in soil soundscapes.

| Index | Aim | Measures | Expectation |
| --- | --- | --- | --- |
| Acoustic Complexity Index (ACI) (Pieretti et al., 2011) | Aims to capture variation in 'complex' biophonic sounds that vary over frequency whilst remaining insensitive to 'simple' anthropophonic or geophonic sounds that stay at a constant frequency such as engine noise. | Calculates the difference in amplitude values between time samples within a frequency band, relative to the total amplitude within that band. | Increasing values in unburnt soils with an increase in biophony. One of a pair of heuristic indices (with BI) intended here to directly reflect biophony. |
| Bioacoustic Index (BI) (Boelman et al., 2008) | Intended as a metric of relative avian abundance on Hawaiian islands, based on both increased amplitude and increased use of frequency bandwidth with increased avian density. | Calculates the area under the curve for frequency spectra at user-specified frequency bands in dB above the minimum dB value. Therefore likely to be influenced by anthropophony and geophony unless carefully filtered recordings are used. | Increasing values with an increase in biophony. One of a pair of heuristic indices (with ACI) intended here to directly reflect biophony. |
| Spectral Events (EVNspCount) (Towsey et al., 2013, QUT, 2023) | A simple measure of acoustic activity in a recording, defined as 'a contiguous set of spectrogram cells/pixels whose decibel values exceed some user defined threshold'. | The number of acoustic events which surpass both an amplitude and duration threshold, calculated from a spectrogram. The values used here are the mean number of events per second over the duration of the 1 minute recording. | Increasing values in unburnt soils with an increase in biophony. One of a pair of indices (with ROIcover) intended here as a metric of general acoustic activity levels. |
| Normalized-Difference Sound Index (NDSI) (Kasten et al., 2012) | Intended as a comparison of the levels of anthropophony and biophony in a soundscape as a metric of human disturbance. Sound below a frequency threshold is considered anthropophony, above it biophony. | Calculated as a ratio of the power spectrum density of the frequencies considered to be anthropophony and biophony. Normally values between -1 (most sound in low frequency band) to +1 (most sound in high frequency band), but we used absolute values as a measure of the even distribution of sound. | Values are expected to be lower in unburnt forest as we expect biophony to be spread over a greater range of frequencies and not solely clustered at very low frequencies. One of a pair of indices (with Hf) intended here as a metric of the distribution of sound across frequency. |
| Regions of Interest (ROIcover) (Ulloa et al., 2021) | A theoretical extension of EVNspCount, looking at the area of a spectrogram that is occupied by sound events, although calculated differently. | Applies an amplitude mask over the spectrogram, regions exceeding the threshold are considered of interest, and the proportion of the spectrogram covered by these regions is returned. | Increasing values in unburnt soils with an increase in biophony. One of a pair of indices with EVNspCount intended here as a metric of general acoustic activity levels. |
| Frequency Entropy (Hf) Sueur et al., (2008) | Intended to measure the dispersion of energy amongst frequency bands. | Calculates the Shannon entropy of a power spectrogram density. Low values reflect sound being clustered along a narrow frequency band. | Values are expected to be higher in unburnt forest as we expect biophony to be spread over a greater range of frequencies and not dominated by a single species at one frequency. One of a pair of indices (with NDSI) intended here as a metric of the distribution of sound across frequency. |

## SOM Appendix 4: Sensitivity analysis of acoustic processing methods

During preliminary investigation it was apparent that strong horizontal banding, likely caused by equipment self-noise, was very prominent in many of the spectrograms between 0-500 Hz, at times appearing to mask biophonic signal. We explored two potential noise reduction methods to increase the prominence of biophonic sounds using the scikit-maad package (Ulloa et al., 2021). Spectral Subtraction (SS) is a common method for the removal of acoustic noise in speech (Boll et al., 1979), implemented using the remove_background() function. The Adaptive Level Equalisation (ALE) algorithm is specifically intended for recordings of the natural environment and it maintains complex acoustic structures such as bird calls (Towsey et al., 2013), this was implemented with the remove_background_along_axis() function. A qualitative assessment of the two denoising techniques over a small subset of the data suggested that SS resulted in reduction but not removal of the bands, whilst the ALE algorithm almost entirely removed the bands, but also removed some of the acoustic signal as well. Consequently we chose to analyse the data using both denoising methods, which resulted in two versions of the dataset.

To test the sensitivity of acoustic indices to the frequency bounds at which they are calculated, we calculated each of the acoustic indices at three different frequency bandwidths, 0-200 Hz, 0-500 Hz and 0-2000 Hz resulting in six sets of values per acoustic index (two denoising methods and three frequency bandwidths). All other parameters for the acoustic indices remained as described in Appendix 3, except for NDSI, with the lower frequency band set at 0 to 100, 150 and 200 Hz respectively and upper frequency band set at 100, 150 and 200 Hz to the maximum of each frequency band respectively. For each of the six sets of index values, and for each of the six indices, we calculate the same generalised linear mixed models as described in the Methods section of the manuscript for the spatial analysis, and compared the results (Figure S4.1).

**Figure S4.1.**
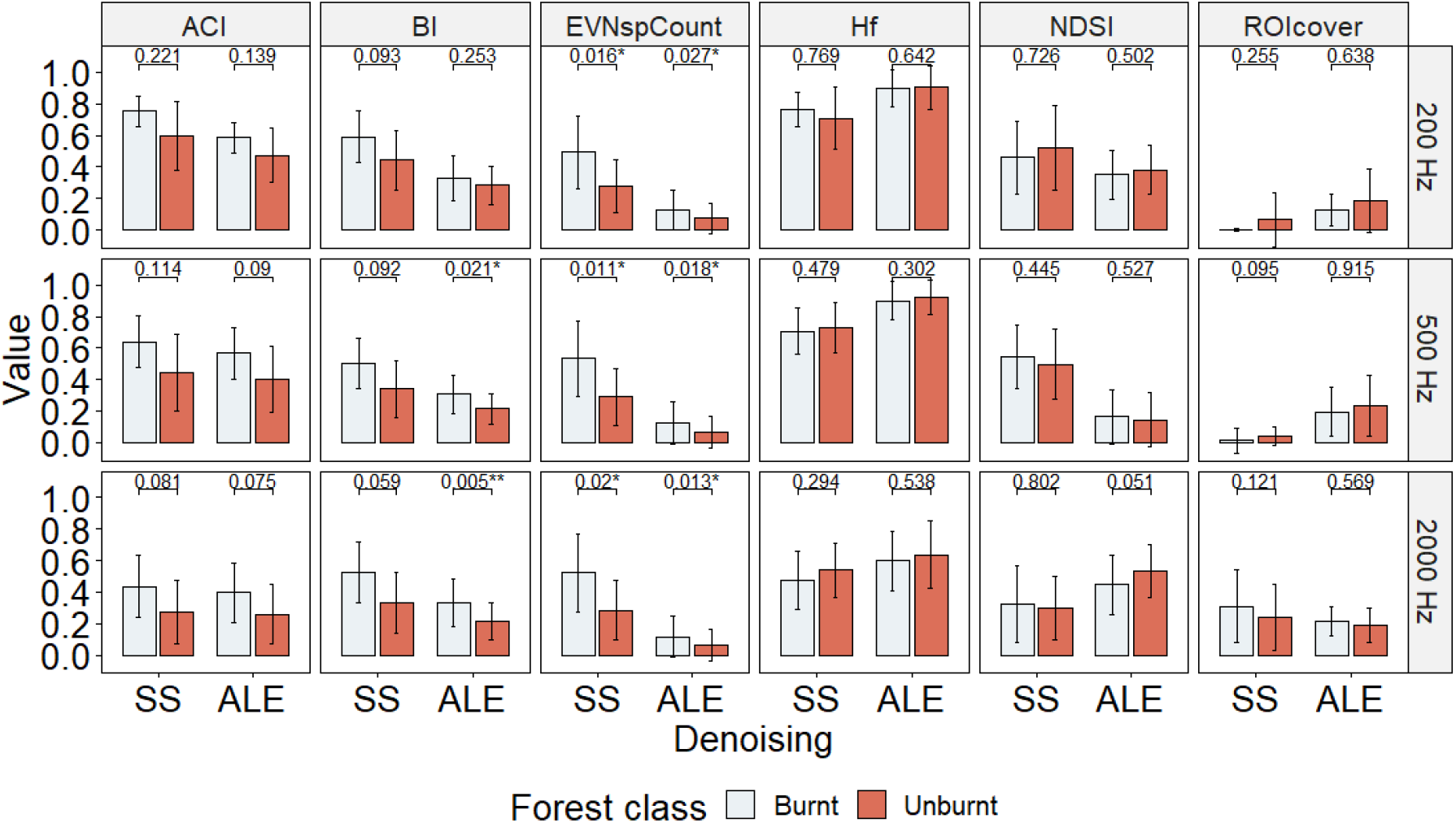
Boxplots of the calculated index values for six acoustic indices, three frequency bands (top 0-200 Hz, middle 0-500 Hz, bottom 0-2000 Hz) and two denoising methods (SS=spectral subtraction, ALE=adaptive level equalisation). Significance values are derived from generalised linear mixed models: ^*^p < 0.05, ^**^ p < 0.01.

### Results and Discussion

Overall, the denoising method only had a limited impact on the soundscape patterns of burnt and unburnt forest. Temporal entropy was the only index that showed any difference in the direction of the relationship between the two methods and then only at a maximum frequency of 200 Hz, when unburnt forest had marginally higher mean values than burnt with ALE, but slightly lower mean values with SS. Otherwise, the impact of denoising method selection was limited to relatively small changes in effect size, consistent with different reductions in the amount of acoustic energy in the spectrogram. Frequency limits in calculation had a slightly bigger impact, for Hf and NDSI unburnt forest had greater mean values at 0-500 Hz and 2000 Hz, but generally lower values at 0-200 Hz, whilst ROIcover had greater values in 0-200 Hz and 0-500 Hz, but lower values in 0-2000 Hz, showing that the frequency chosen to calculate indices can impact results.

